# Simultaneous production of diverse neuronal subtypes during early corticogenesis

**DOI:** 10.1101/369678

**Authors:** E. Magrinelli, R. J. Wagener, D. Jabaudon

## Abstract

The circuits of the neocortex are composed of a broad diversity of neuronal cell types, which can be distinguished by their laminar location, molecular identity, and connectivity. During embryogenesis, successive generations of glutamatergic neurons are sequentially born from progenitors located in germinal zones below the cortex. In this process, the earliest-born generations of neurons differentiate to reside in deep layers, while later-born daughter neurons reside in more superficial layers. Although the aggregate competence of progenitors to produce successive subtypes of neurons progresses as corticogenesis proceeds, a fine-grained temporal understanding of how neuronal subtypes are sequentially produced is still missing. Here, we use FlashTag, a high temporal resolution labeling approach, to follow the fate of the simultaneously-born daughter neurons of ventricular zone progenitors at multiple stages of corticogenesis. Our findings reveal a bimodal regulation in the diversity of neurons being produced at single time points of corticogenesis. Initially, distinct subtypes of deep-layer neurons are simultaneously produced, as defined by their laminar location, molecular identity and connectivity. Later on, instead, instantaneous neuronal production is homogeneous and the distinct superficial-layer neurons subtypes are sequentially produced. These findings suggest that early-born, deep-layer neurons have a less determined fate potential than later-born superficial layer neurons, which may reflect the progressive implementation of pre-and/or post-mitotic mechanisms controlling neuronal fate reliability.

## Introduction

The neocortex is a six-layered structure containing a large diversity of neuronal cell types, which can be defined by the combination of their laminar position, molecular identity and connectivity (Harris and Shepherd 2015; Tasic et al. 2016; Jabaudon 2017). From an ontological and developmental perspective, glutamatergic cortical neurons can be divided into two main classes: deep-layer neurons (*i.e.* neurons which are located in layer (L)6 and L5), which are predominantly subcortically-projecting, of which about 18 transcriptionally distinct subtypes have recently been described in the visual cortex, and superficial layer neurons (L4 and L2/3), which are predominantly intracortically-projecting, of which 5 transcriptional subtypes have been described (Florio et al. 2014; Lui et al. 2011; Govindan and Jabaudon 2017; Tasic et al. 2017). Glutamatergic neurons are born from progenitors located below the developing neocortex, from where they migrate radially to their final laminar destination and differentiate into mature neuronal subtypes. Cortical neurons are born either directly from apical progenitors (APs) located in the ventricular zone (which lines the ventricles), or indirectly via intermediate progenitors, located in the subventricular zone above it (Lui et al. 2011; Govindan and Jabaudon 2017). During corticogenesis, these progenitors first generate deep-layer neurons, followed by superficial-layer neurons, in a so-called inside-out process of neuronal production (Kornack and Rakic 1995; Angevine and Sidman 1961; Polleux et al. 1997; Takahashi et al. 1999).

While the overall sequential generation of deep- and superficial-layer neurons is well established, our understanding of this process at high temporal resolution is still partial: at a given timepoint in corticogenesis, are single neuronal subtypes produced, or is there heterogeneity within the successive generations of daughter neurons? In particular, given that in mice a roughly equal amount of time is devoted to the generation of deep-layer vs. superficial-layer neurons (3-4 days) despite a seemingly broader diversity of molecular subtypes of deep-layer neurons (Tasic et al. 2016, 2017), could distinct subtypes be simultaneously generated early on?

This question has been difficult to address using traditional birth dating approaches such as thymidine analogue pulse-labeling, since this method labels progenitors over the several hours of duration of the S phase or, when using combinatorial S-phased based approaches to increase temporal resolution, is not selective for APs vs. BPs (see *e.g.* Takahashi, et al. 1996, and review in Govindan et al., 2018). This represents obstacles to link time of birth with final fate. To circumvent these limitations, we recently developed the FlashTag approach, which allows specific labeling of APs precisely at the time neurons are born, by taking advantage of the fact that these cells divide in contact with the ventricular wall (Telley et al. 2016; Govindan et al. 2018). This allows for high temporal resolution tagging of simultaneously-born generations of AP daughter neurons, which can be tracked as they migrate and differentiate.

Using FlashTag, we reveal a dynamic regulation in the diversity of neurons that are simultaneously produced by APs at single time points of development. At early stages of corticogenesis, as deep-layer neurons are being generated, we observe an unexpected diversity in the final identities of simultaneously-born neurons, as assessed by a variety of laminar, molecular and connectivity features. Later in corticogenesis, instead, APs give birth to neurons with more homogenous features, which tightly correspond to their time of birth. Together, these findings reveal an initially broad neuronal fate diversity early in corticogenesis, which narrows down as corticogenesis proceeds, suggesting the progressive implementation of mechanisms controlling neuronal production and/or differentiation.

## Results

We used FlashTag (FT) pulse-labeling to determine the fate of synchronously-born cohorts of neurons on sequential embryonic days (E) between E11.5 and E16.5 in the mouse primary somatosensory cortex. This period includes the time of generation of deep-layer neurons (E11.5-E13.5) and superficial-layer neurons (E14.5-E16.5). FT^+^ neurons overwhelmingly correspond to AP-born daughter cells (Telley et al. 2016; Govindan et al. 2018); here, to allow for even better temporal resolution, in most experiments we combined this approach with the chronic delivery of BrdU via an intraperitoneal osmotic pump. Using this approach, neurons born directly from APs can be identified as FT^+^ BrdU^-^ cells, whereas neurons which have undergone intercurrent divisions are BrdU^+^ (Fig. 1a and Fig. S1a; Telley et al., 2016; Govindan et al., 2018).

**Figure 1.**
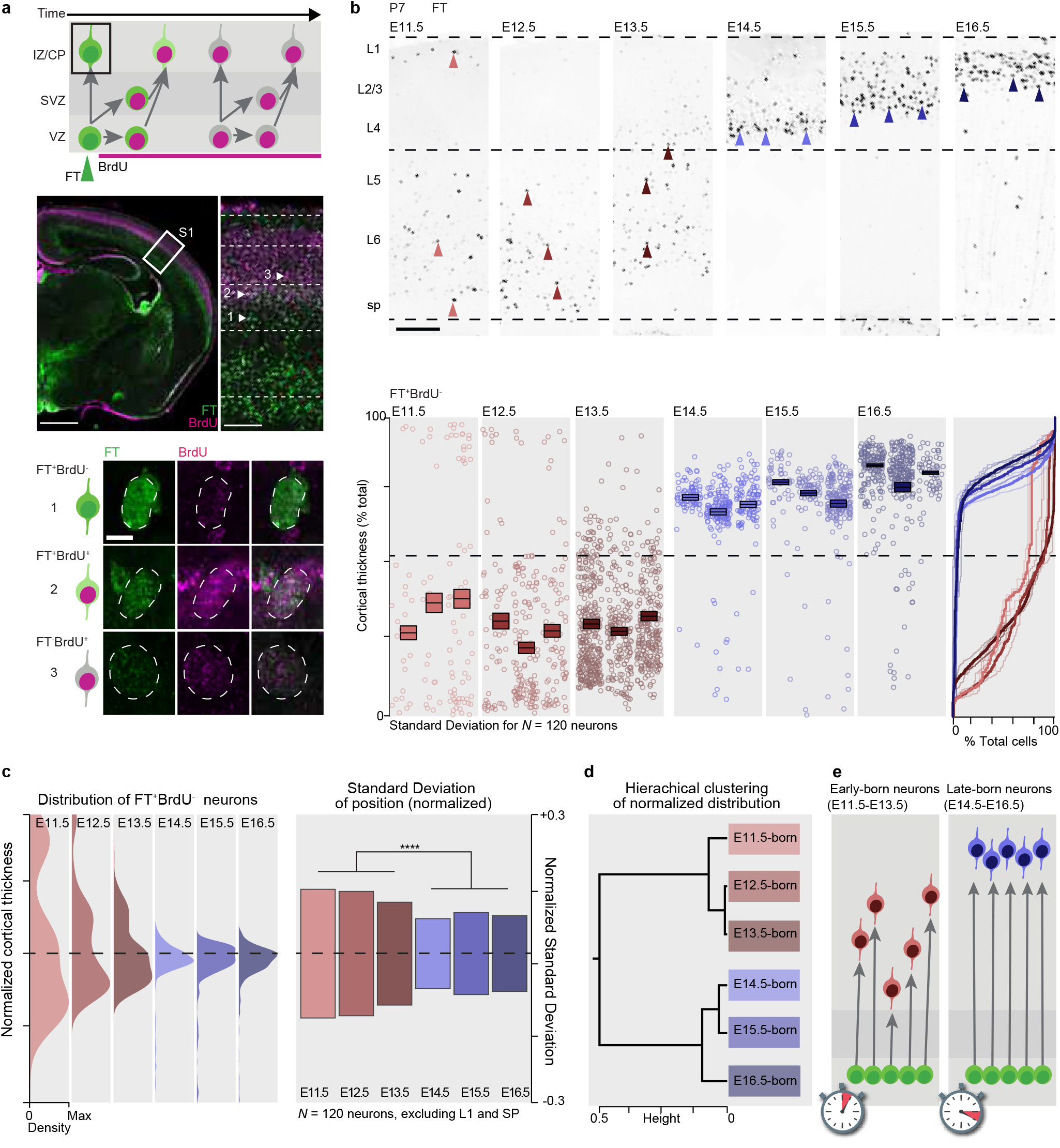
Simultaneously-born, deep-layer neurons have a broad range of laminar fates. **a,** Top: Schematic representation of the FlashTag (FT) labeling strategy. Bottom: Photomicrographs illustrating the results of FT-chronic BrdU labeling, highlighting FT^+^ BrdU^-^ (*i.e.* directly AP-born) neurons, which are studied here. **b,** P7 laminar distribution of AP-born neurons at different embryonic ages. Top: Representative photomicrographs; bottom: Quantifications. **c,** Left: density plot of mean-normalized neuronal distribution. Right: Mean-normalized standard deviation (**** *P* = 0.0008, unpaired T-test between E11.5-E13.5-born and E14.5-E16.5-born neurons). **d**, Unsupervised hierarchical clustering of radial position at each birth time. **e,** Summary of the findings. Scale bars: 1mm (a, left), 200 μm (a, right), 5 μm (a, bottom), 120μm (b). CP – Cortical Plate, E – Embryonic day, IZ – Intermediate Zone, L – Layer, P – Postnatal day, RGC – Radial Glia Cell, SVZ – Subventricular Zone, S1 – Primary somatosensory cortex, VZ – Ventricular Zone.

We first focused on the laminar fate of synchronously-born neurons at these sequential embryonic stages by assessing the radial position of labelled neurons at P7, once migration is complete (Fig. 1b and Fig S1a). We observed that neurons born at early stages of corticogenesis (E11.5-E13.5) distribute broadly within deep cortical layers, whereas at later stages (E14.5-E16.5) neurons are laminarly compact and their radial location closely corresponds to time of birth (Fig. 1b-e). This difference persisted even when not considering L1 and subplate neurons (which are also generated at E11.5 and E12.5), and thus corresponded to a genuine diversity in distribution within deep layers (Fig. 1c). Confirming this bimodal, birthdate-dependent diversity, unbiased cluster analysis of the normalized standard deviation in final radial position (calculated using 120 randomly-selected neurons at all stages, see Methods) revealed that E11.5- E13.5-born neurons on the one hand, and E14.5-E16.5-born neurons on the other hand, have different distribution patterns (Fig. 1d).

The data above reveal that time of birth becomes a stronger determinant of final radial location later in corticogenesis. This suggests that sequential generations of neurons born at a fixed interval may have overlapping laminar locations if born early, but distinct distributions if born late. To test this possibility, we labelled two sequentially-generated cohorts of neurons with two pulses of FT administered at a 6-hour interval in single embryos (Fig. 2). Distinct dye colors were used in order to distinguish the laminar location of the two AP-born daughter neuron cohorts at P7. Confirming our hypothesis, successively-born neurons had overlapping distributions when born at E13.5 and E13.5 + 6 hours (Fig. 2a, *P* = 0.34, paired t-test), whereas E15.5- and E15.5 + 6 hour-born neurons had distinct laminar identities (Fig. 2b, *P* < 10^-4^, paired t-test). Together, these data reveal that during early corticogenesis, sequentially-born neurons have overlapping laminar fates, whereas later on, neurons with a similarly interval in birthdate have distinct laminar fates.

**Figure 2.**
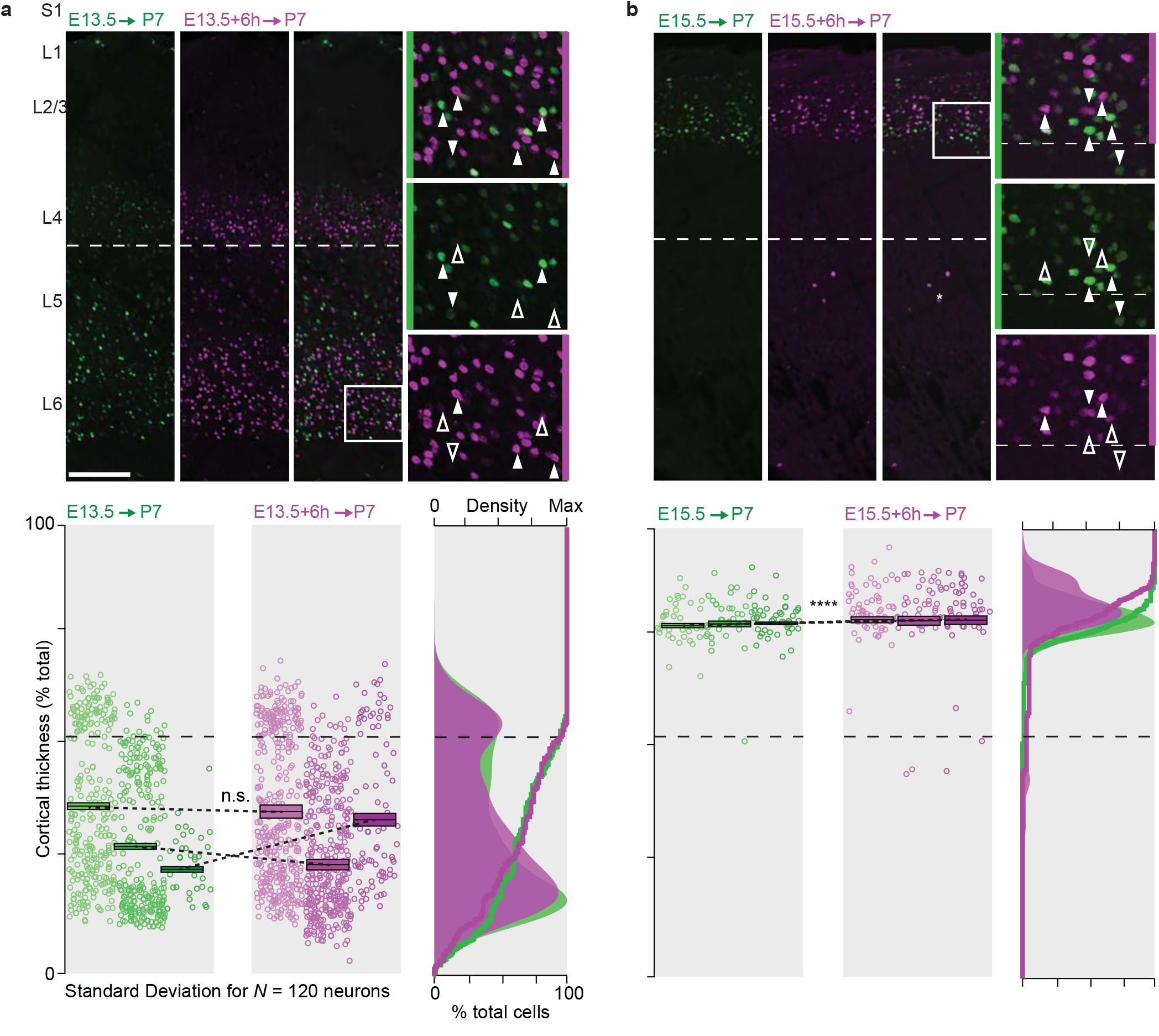
Closely sequentially-born neurons have overlapping distribution early but not late in corticogenesis. **a**, Neurons born at E13.5, and E13.5 + 6h weighted paired t-test have overlapping laminar locations at P7 (*P* = 0.34, paired t-test). **b**, Neurons born at E15.5, and E15.5 + 6h weighted paired t-test have distinct laminar locations at P7 (*P* < 10^-4^, paired t-test). Scale bars: 100μm.

One explanation for the greater laminar dispersion of early-born neurons could be that deep-layer neurons are initially laminarly compact, but are subsequently dispersed by the migration of successive waves of later-born neurons. We thus examined when during neuronal differentiation heterogeneities within simultaneously-generated cohorts of neurons first appear. For this purpose, we FT pulse-labelled neurons at either E13.5 (to follow of deep-layer neurons) or E15.5 (to follow superficial-layer neurons) and tracked the radial location of FT^+^ cells at 24 hours intervals throughout corticogenesis (Fig. 3a and b). E13.5-born neurons reached the cortical plate within 48 hours after their birth, and their distribution within deep cortical layers was broad from the onset on, *i.e.* we did not observe a progressive dispersion of these cells over time (Fig. 3c; *P* = 0.41, one-way ANOVA). E15.5- born neurons took an additional day to reach the cortex (*i.e.* a total of 72 hours) and rapidly aligned to form a compact homogeneous layer (Fig. 3c; *P* = 0.03, one-way ANOVA). Together, these data indicate that the overlapping distribution of sequential generations of early-born neurons within deep cortical layers is not secondary to the subsequent arrival of later-born neurons, but instead is the direct consequence of migration to a broader diversity of laminar targets.

**Figure 3.**
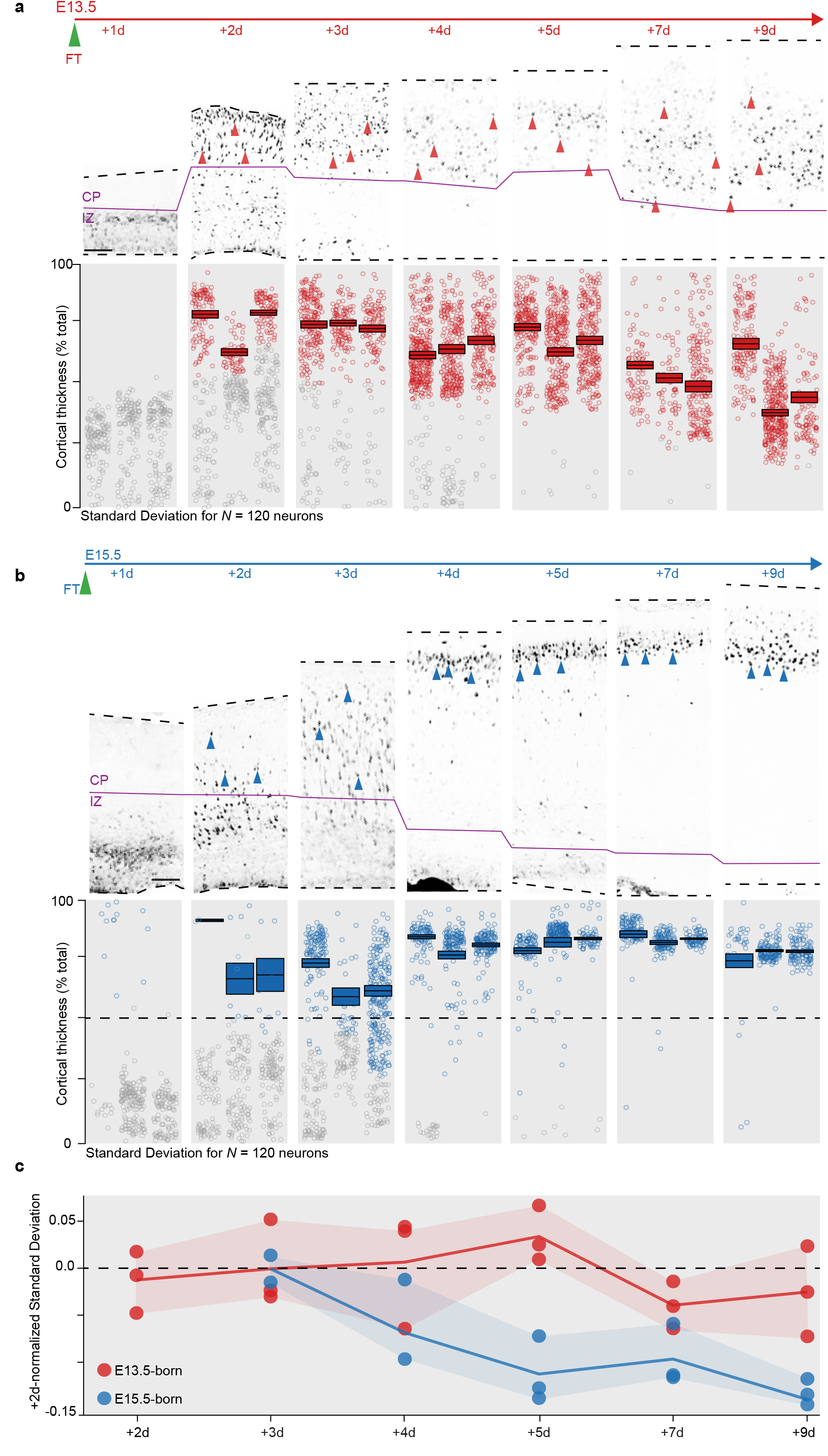
Early-born neurons are dispersed as soon as they invade the cortical plate. **a**, Migration dynamics of simultaneously-born E13.5 neurons from 1 day to 9 days after FT injection. **b**, Migration dynamics of simultaneously-born E15.5 neurons from 1 day to 9 days after FT injection. Note the rapidly compact laminar distribution of these neurons compared to a. **c**, Standard deviation of position in the cortical plate for E13.5- and E15.5-born neurons. The standard deviation for E15.5-born neurons rapidly decreases while it remains essentially constant for s E13.5-born neurons. Scale bars: 50μm (a, b).

We next investigated whether the laminar diversity in simultaneously-produced E13.5-born neurons was accompanied by a corresponding diversity in their molecular identity. To this end, we examined the expression of deep- (TBR1, CTIP2) and superficial- (SATB2, CUX1) layer neuronal markers in birthdate-identified cells at P7 (Fig. 4, Figs. S2 and S3. Hevner et al. 2002; Arlotta et al. 2005; Alcamo et al. 2008; Rodríguez-Tornos et al. 2016). This approach revealed that simultaneously-produced E13.5-born neurons are molecularly heterogeneous, since only a small fraction of these cells expressed each of these four makers, which they progressively switch on during their differentiation (Fig. S4). Importantly, molecular expression was congruent with the laminar position of these cells. For example, the L5 marker CTIP2 was essentially exclusively expressed by neurons located in L5, and the superficial layer marker CUX1 was predominantly expressed by the few neurons present in L4. In contrast, protein expression was much more homogeneous in E15.5-born neurons, consistent with the convergent laminar identity of these cells (Fig. S3). Thus, laminarly and molecularly diverse neurons are simultaneously generated at early stages of corticogenesis, whereas neuronal production becomes more homogeneous later on.

**Figure 4.**
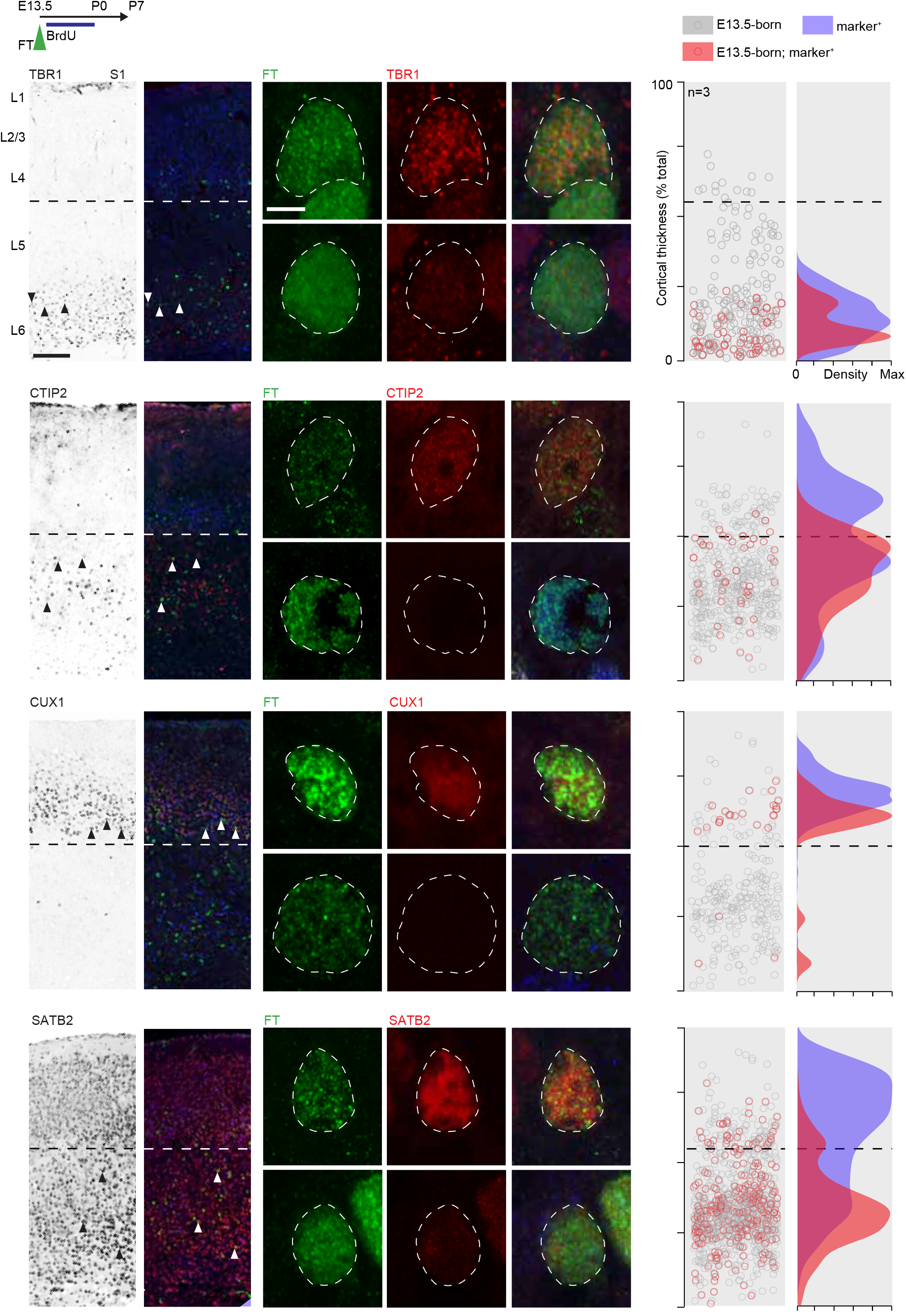
Simultaneously-born deep-layer neurons have diverse molecular identities. Immunostaining of E13.5 AP-born neurons with TBR1, CTIP2, CUX1, and SATB2 antibodies reveals distinct molecular identities, which match their laminar locations. Photomicrographs in first column: antibody alone, second column: antibody + FT, following columns: high magnifications. Scale bars: 200μm (low magnification), 4μm (high magnification).

Finally, we assessed the connectivity of these laminarly and molecularly distinct subtypes of simultaneously produced, deep-layer neurons. Using retrograde labeling from distinct subcortical (thalamus and spinal cord) and intracortical (contralateral hemisphere) targets, we observed that although simultaneously born, these neurons have a diversity of axonal projections, which corresponds to their laminar and molecular identity (Fig. 5, S5). For example, corticothalamic projection neurons were located in L6, whereas corticospinal neurons were confined to L5. Thus, the laminar and molecular diversity of simultaneously-produced deep-layer neurons is accompanied by a corresponding diversity in the connectivity of these cells.

**Figure 5.**
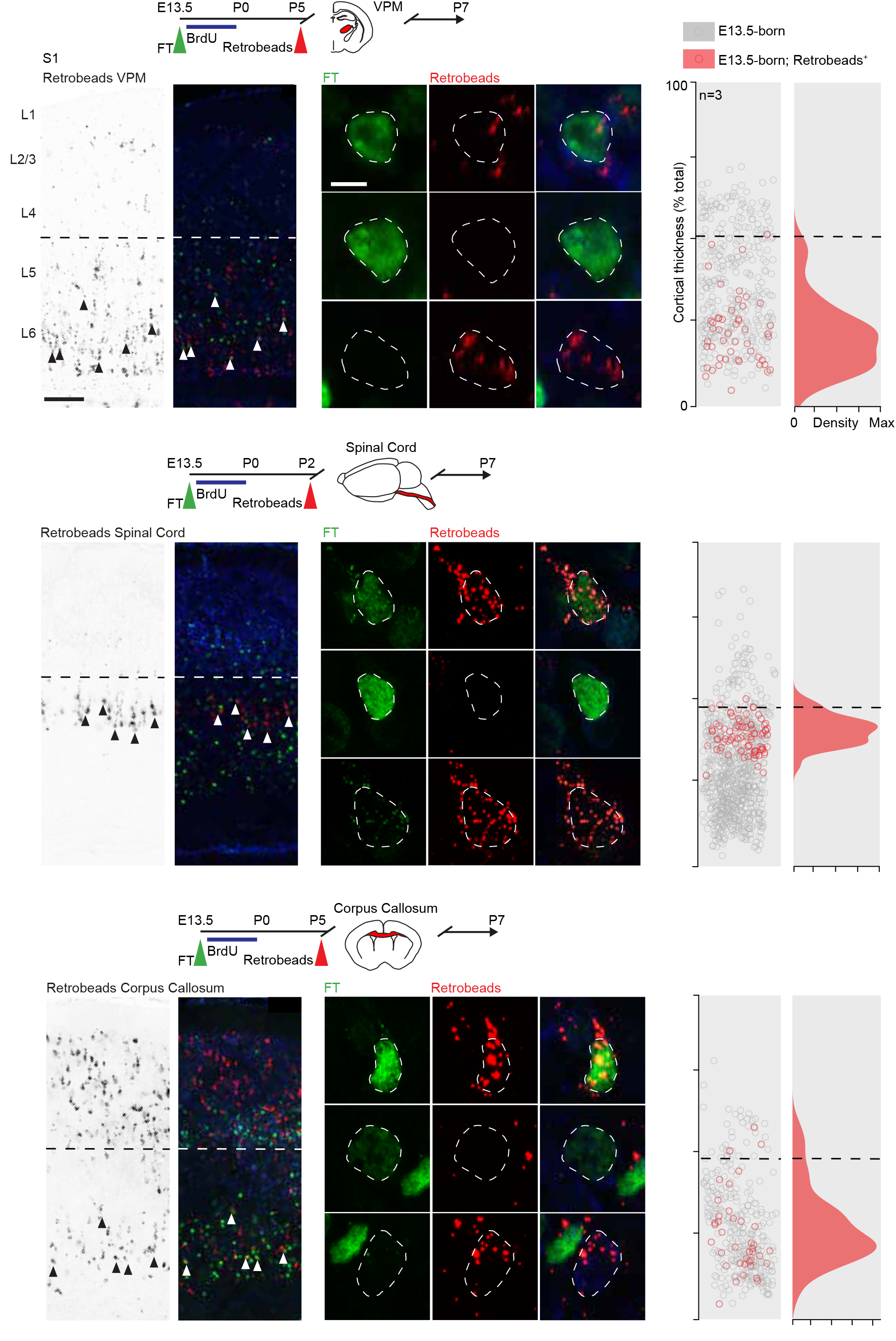
Simultaneously-born deep-layer neurons have diverse connectivities. Retrograde labeling of E13.5 simultaneously-born neurons reveals a diversity of projections to the ventroposterior medial nucleus of the thalamus (VPM), spinal cord and corpus callosum. Photomicrographs in first column: retrograde labeling alone, second column: retrograde labeling + FT, following columns: high magnifications. Scale bars: 200μm (low magnification), 5μm (high magnification).

## Discussion

Our findings reveal that early in corticogenesis, distinct subtypes of deep-layer neurons are simultaneously produced from APs, whereas later on, the distinct superficial layer neuron subtypes are sequentially born. Before E14.5, the correlation between time of birth and neuronal identity is thus initially relatively loose and tightens later on as superficial layer, intracortically-projecting neurons are being born. These findings unveil a dynamic regulation over neuronal fate choice and suggest the progressive implementation of pre- and/or post-mitotic mechanisms controlling the acquisition of final neuronal identity. The simultaneous production of neurons with distinct laminar fates has been reported before, and is particularly visible in species with prolonged corticogenesis, such as primates (Rakic 1982; Granger et al. 1995; Takahashi et al. 1999); these findings imply some level of heterogeneity in progenitors or in the differentiation paths of daughter neurons. For example, the simultaneous production of neurons born directly from APs or indirectly *via* intermediate progenitors (which are not distinguished by traditional thymidine analogue birth dating studies) may generate neurons with distinct final properties (Tyler et al. 2015; Guillamon-Vivancos et al. 2018). Here in addition, using FlashTag birthdating (Govindan et al. 2018), we reveal that AP-born daughter neurons born simultaneously have different fates if generated early in corticogenesis.

One possible scenario accounting for a temporally-regulated, pre-mitotic control over daughter neuron fate is that diverse, fate-restricted progenitors are present early on, while a more homogeneous population of progenitors gives rise to superficial layer neurons. Although the existence of fate-restricted progenitors has been proposed to account for the generation of deep- vs. superficial-layer neurons (Frantz and McConnell 1996; Franco et al. 2012; Costa and Müller 2014; Gil-Sanz et al. 2015), there is otherwise little evidence for subtype-specific progenitors, even using modern single-cell RNA sequencing approaches (Guo et al. 2013; Yuzwa et al. 2017). Another possibility is that early progenitors are permissive (as opposed to instructive) for the emergence of neurons with multiple potential fates. This could for instance occur if early progenitor chromatin structure offers the possibility for daughter neurons to differentiate along multiple (even simultaneous) paths and be refined post-mitotically (see below). Of note, a permissive chromatin state may be required for the progression in the neurogenic competence of early APs, which may be less likely the case for later-born APs, whose natural neurogenic competence becomes more restricted. The progressive implementation of epigenetic control mechanisms may thus allow for the generation of more reliable, albeit less diverse, neuronal cell types (Nitarska et al. 2016; Bonev et al. 2017; de la Torre-Ubieta et al. 2018). In a similar line of thought, cell cycle length progressively increases during corticogenesis (Calegari et al. 2005; Borrell and Calegari 2014; Dehay and Kennedy 2007), perhaps allowing more time for “quality control” at later developmental stages.

With regard to post-mitotic control over daughter neuron fate, acquisition of final identity may be more strongly dependent on post-mitotic events for early-born neurons. Thus, diversity in final identity might be secondary to select migration events and interactions with the environment. Deep- and superficial-layer neurons evolve in distinct environments when migrating from their place of birth to their final laminar target, and they do so with different kinetics and using distinct strategies. For example, at later developmental stages, migrating neurons undergo a stalling period in the subventricular zone before entering the cortical plate (Ohtaka-Maruyama et al. 2018), which may play a role in coordinating laminar (and overall) neuronal identity. In addition, interactions with subplate neurons (Ozair et al. 2018; Ohtaka-Maruyama et al. 2018) and a potential non-cell autonomous role for laminar position in the acquisition of final identity (Oishi et al. 2016) may create a context in which final specification is more strongly dependent on (potentially less deterministic) post-mitotic events for early-born neurons than for later-born neurons.

In evolutionary terms, deep-layer neurons have been thought to be phylogenetically older than superficial layer neurons (Aboitiz and Zamorano 2013; Jabaudon 2017; but see Tosches et al. 2018). In this perspective, superficial layer neurons represent a specialized, evolutionary novel acquisition, over the development of which several levels of molecular controls may have been superimposed. In contrast, in “primordial” deep-layer neurons, increased variability may have been selected over reliability in order to generate diverse neuronal cell types in a limited amount of time. The dynamic regulation over the generation of neuronal diversity identified here may thus represent an evolutionary comprise to allow both reliable and diverse circuits to develop in expanding mammalian brains.

## Acknowledgments

We thank the Imaging Platform of the University of Geneva, A. Benoit for technical assistance; and members of the Jabaudon laboratory for constructive comments on the manuscript. The Jabaudon laboratory is supported by the Swiss National Science Foundation; E.M. is supported by EMBO Long Term Fellowship.

## Author contributions

E.M., and D.J. conceived the project and designed the experiments. E.M., and R. J. W. performed the experiments. E.M. and D.J. wrote the manuscript.

## Methods

### Mice

CD1 male and female mice were used for all experiments and with the permission of the Geneva Cantonal Authorities, Switzerland. Matings were performed over a 3-hour window, which was considered as time E0.

### In utero FlashTag injections

Pregnant mice were anesthetized by isoflurane at precise gestation time points and placed on a warm operating table. Small abdominal incisions were performed to expose uterine horns, and FlashTag was injected in the ventricles (see Govindan et. al 2018 for details). The double Flash Tag experiment was carried out injecting CFSE (carboxyfluorescein succinimidyl ester; CellTrace^™^ Life Technologies, #C34554) and CellTrace Violet (Life Technologies, #C34557). 414 nl of FlashTag was injected in the third ventricle at embryonic ages E11.5-E15.5, allowing for diffusion in lateral ventricles. For E16.5 embryos injections were performed directly in the lateral ventricle using 207nl of FlashTag. At the end of the procedure, the uterine horns were reintroduced in the abdominal cavity and peritoneum and skin were independently sutured. Mice were kept on a heating pad until recovery from the anaesthesia.

### Chronic BrdU delivery

Chronic BrdU delivery was performed using osmotic pumps loaded fully with 16mg/ml BrdU in PBS. 3 days 0.1ul per hour (1003D Alzet) and 7 days 0.1ul per hour (2001 Alzet) osmotic pumps were used to cover the necessary delivery period. Osmotic pumps were introduced in the peritoneal cavity while performing in utero injections.

### Retrograde labeling

Callosal and thalamic (ventroposterior medial nucleus, VPM) injections were performed using stereotaxic guided injection. P5 pups were anesthetized on ice. Heads were fixed in a Digital Lab Standard Stereotaxic Instrument (Stoelting 51900). A small incision was performed on the top of the skull to visualize the bregma for stereotaxic references. For retrograde labeling, red Retrobeads^™^ (Lumafluor, Inc.) were loaded in a glass capillary mounted on a Nanoinjector (Nanoject II Auto-Nanoliter Injector, Drummond Scientific Company 3-000-204) and injected as 10 x 18nl injections. Corpus callosum coordinates were (from bregma); X: 0.0, Y: +0.3, Z: −1.4; VPM coordinates were X: −1.2, Y: −0.9, Z: −2.5. Spinal cord injections were performed under ultrasound guided injections using a Vevo 770 ultrasound backscatter microscopy system (Visual Sonics, Canada).

### Post Mortem tissue collection

Embryonic tissue was collected by microdissection in ice-cold PBS. Embryonic brains were fixed in paraformaldehyde (PFA) 4% overnight at +4°C. Postnatal brains were extracted after intracardiac perfusion of PBS and PFA 4% under thiopental anaesthesia and subsequently fixed overnight in PFA 4% at +4°C. Brain samples younger than E15.5 were embedded in 4% selec-agar PBS and cut on a Leica vibrating microtome (Leica, #VT100S) in 50 um-thick coronal free-floating slices. Brains older than E15.5 were equilibrated in sucrose 30% PBS, embedded in Optimal Cutting Temperature (OCT) medium (JUNG, Germany) and cut on a Leica cryostat into 60um coronal free-floating slices.

### Immunohistochemistry and Imaging

All free-floating sections, were washed three times 10 minutes in PBS, incubated one hour at room temperature in blocking solution (4% Bovine Albumin Serum, 0.2% Triton-X 100 in PBS) and then incubated overnight at +4°C with primary antibody diluted in the same blocking solution. Sections were then washed three times in PBS for 15 minutes and incubated 2h with secondary antibodies diluted in blocking solution. After washing again three times 15 minutes with PBS, sections were mounted in Sigma Fluoromount (#F4680). For BrdU antibody staining, sections were denaturated before blocking by incubating them in 2N HCl at 37°C for 40 minutes and washed three times in PBS 15 minutes before blocking. For all experiments using both anti-BrdU and anti-Satb2 primary antibodies, a first overnight primary antibody incubation with only anti-BrdU was performed and then, after washing three times 15 minutes in PBS, a second overnight primary overnight antibody incubation with all remaining antibodies was done, in order to prevent a cross-reaction between the anti-BrdU and anti-Satb2 antibody.

#### Antibodies

Rat anti-BrdU (1:250, Abcam, #AB6326), rat anti-CTIP2 (1:500, Abcam, #AB18465), rabbit anti-CTIP2 (1:500, Abcam, #AB28448), rabbit anti-CUX1 (1:500, Santa Cruz, #sc-13024), rabbit anti-FITC (1:2000, Abcam, #AB19491), goat anti-FITC (1:1000, Novus Biolab, #NB600-493), mouse anti-SATB2 (1:200, Abcam, #AB51502), rabbit anti-TBR1 (1:500, Abcam, #AB31940).

#### Imaging

Images were obtained using either a Nikon A1R spectral confocal and ZEISS LSM 800 for confocal acquisitions, from the University of Geneva bioimaging facility, while epifluorescence images were obtained on an Eclipse 90i Nikon epifluorescence microscope.

### Quantifications

All images quantifications were performed using standard Fiji functionalities. Radial position was measured by recording coordinates of manually-counted cells. To compare radial position between animals, the thickness of the cortex was normalized across animals, as was the thickness of CP for samples of brains before P7. All colocalization analysis were performed by manual analysis of confocal images. In Fig 1c, Unpaired test analysis was performed between E11.5, E12.5, E13.5 injected brains against E14,5, E15,5 and E16,5 injected ones. In Fig. 1d, hierarchical clustering was performed with Euclidean distance calculation. Normalized density distribution was performed with radial position normalized to the mean per age. Paired test analysis in Fig. 2b, was performed between E13.5, E13.5 + 6h, E15.5-P7 and E15.5 + 6h cells, randomly sampling 120 cells for each of the images analyzed. One-way ANOVA analysis of Fig. 3c, was performed for E13.5 and E15.5-born neurons across timepoints, randomly sampling 120 cells for each of the images analyzed. For Fig. S1b, ROI were determined by automatic threshold segmentation with Fiji, including the lowest FT intensity labelled cells. For each ROI the mean of FT signal was recorded as well as the positive or negative BrdU labeling. All data analysis scripts were custom-prepared in R.

---

**Figure S1.**
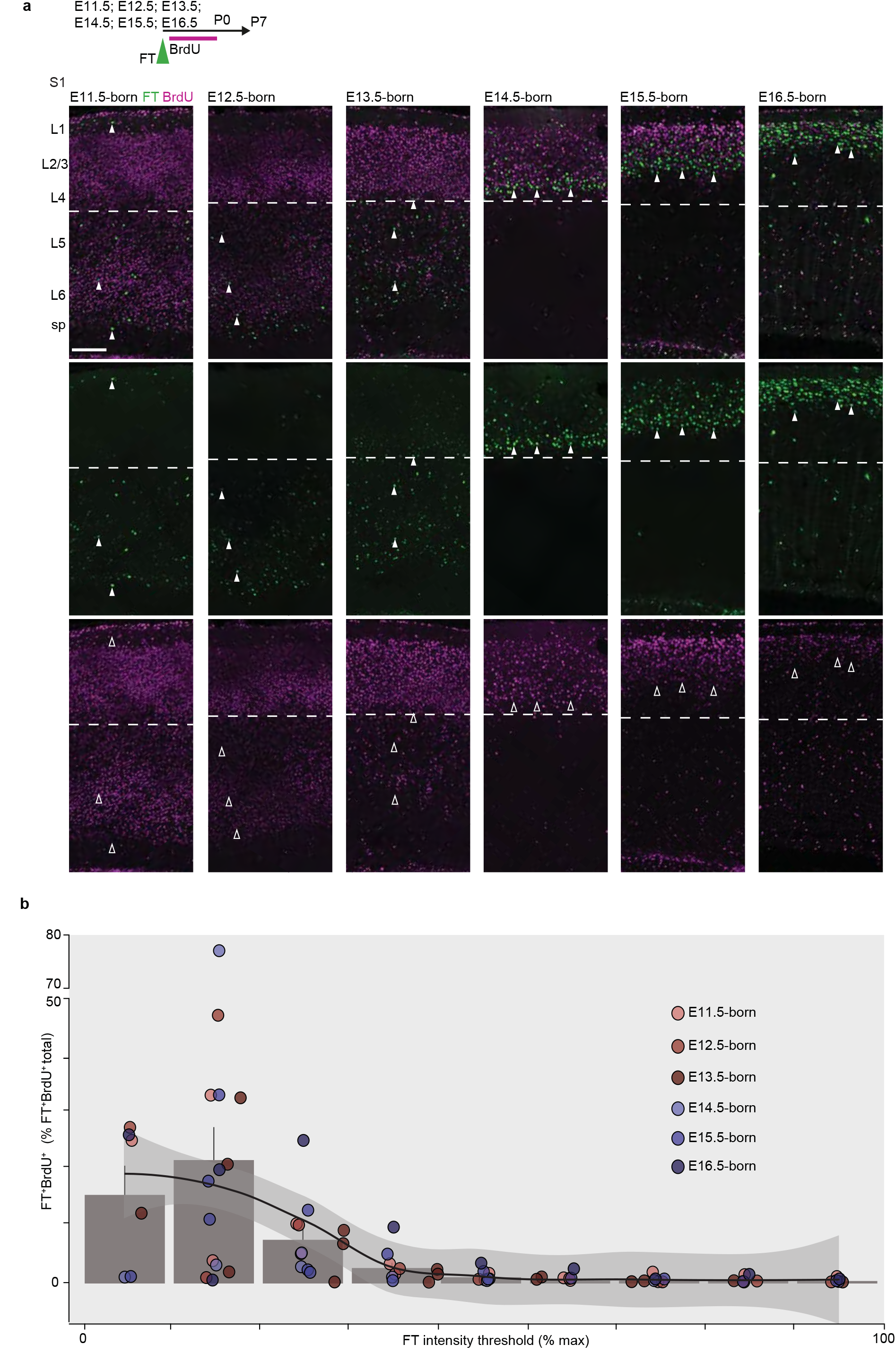
Diversity of laminar contribution of direct neurogenesis across corticogenesis. **a**, Immunostaining for FT and chronic BrdU injected from E11.5 to E16.5. **b**, High FT signal thresholding allows to select for FT^+^;BrdU^-^ neurons, justifying the use of top 10% FT signal as a way to detect directly born neurons without chronic BrdU. Continuous line shows automatic chosen loess interpolation model. Scale bar: 120μm. E – embryonic day, FT – FlashTag, L – Layer, P – Postnatal day, S1 – Primary Somatosensory area.

**Figure S2.**
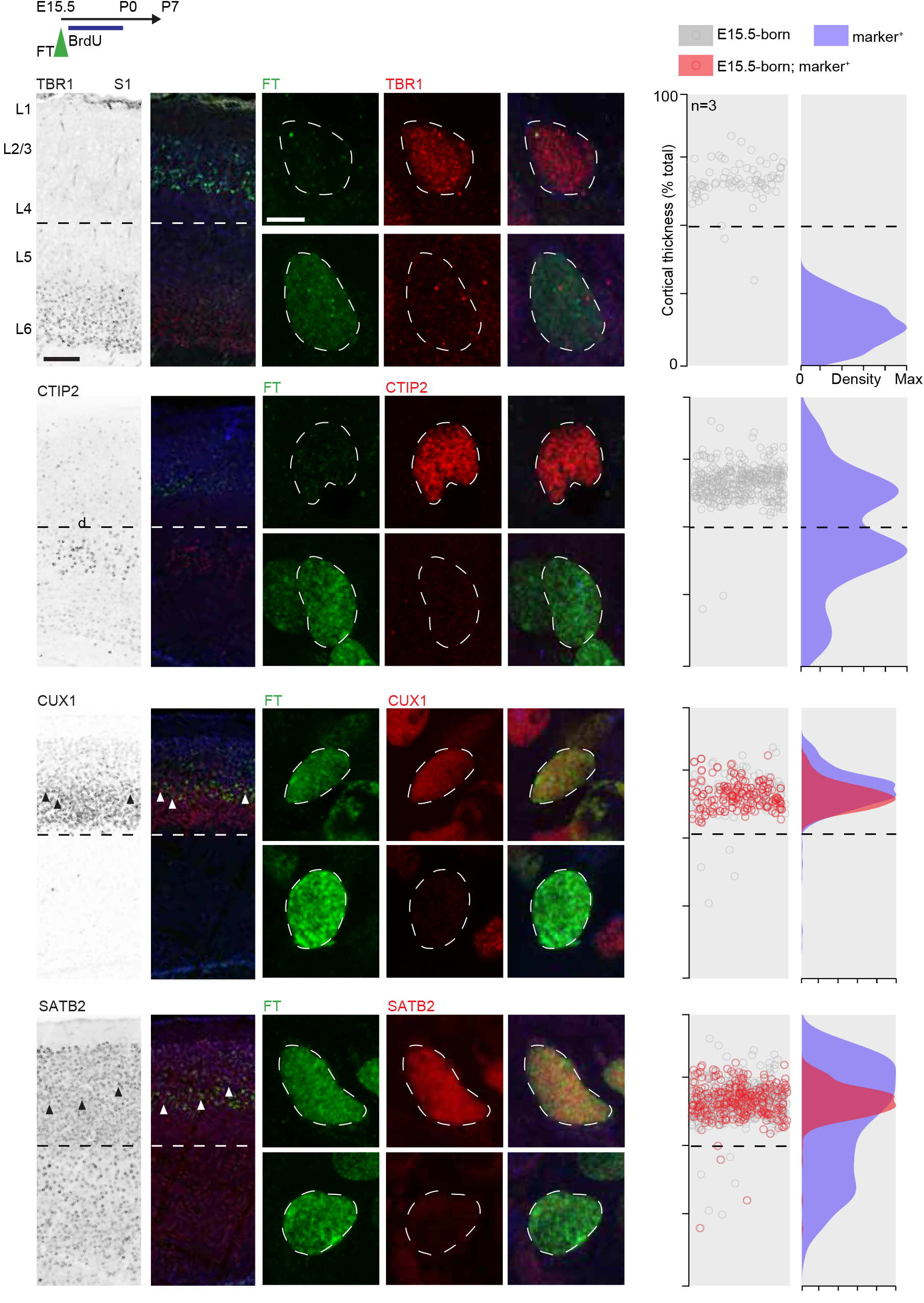
Simultaneously-born superficial-layer neurons have less diverse molecular identity compared to deep-layer ones. Immunostaining of E15.5 AP-born neurons with TBR1, CTIP2, CUX1, and SATB2 antibodies reveals homogeneous molecular identities, which are in agreement with their laminar locations. Scale bars: 200μm (low mag), 4μm (high mag). E – embryonic day, FT – FlashTag, L – Layer, P – Postnatal day, S1 – Primary Somatosensory area.

**Figure S3.**
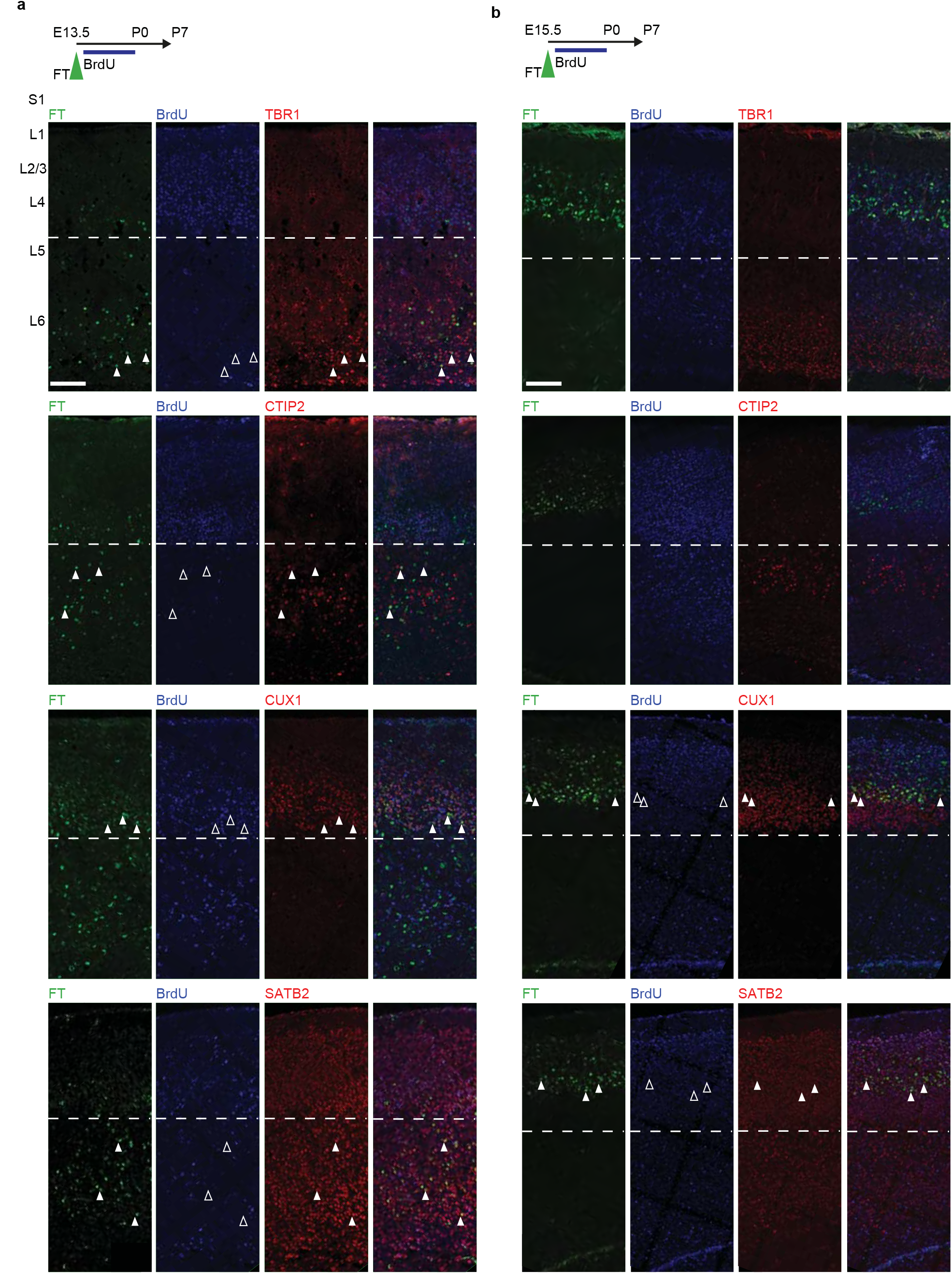
Laminar and molecular diversity of early and late simultaneously-born neurons. **a**, Immunostaining of E13.5 AP-born neurons with TBR1, CTIP2, CUX1, and SATB2 antibodies reveals distinct molecular identities, which are in agreement with their laminar locations. **b**, Immunostaining of E15.5 AP-born neurons with TBR1, CTIP2, CUX1, and SATB2 antibodies reveals homogeneous molecular identities, which are in agreement with their laminar locations. Scale bars: 200μm (a, b). E – embryonic day, FT – FlashTag, L – Layer, P – Postnatal day, S1 – primary somatosensory area.

**Figure S4.**
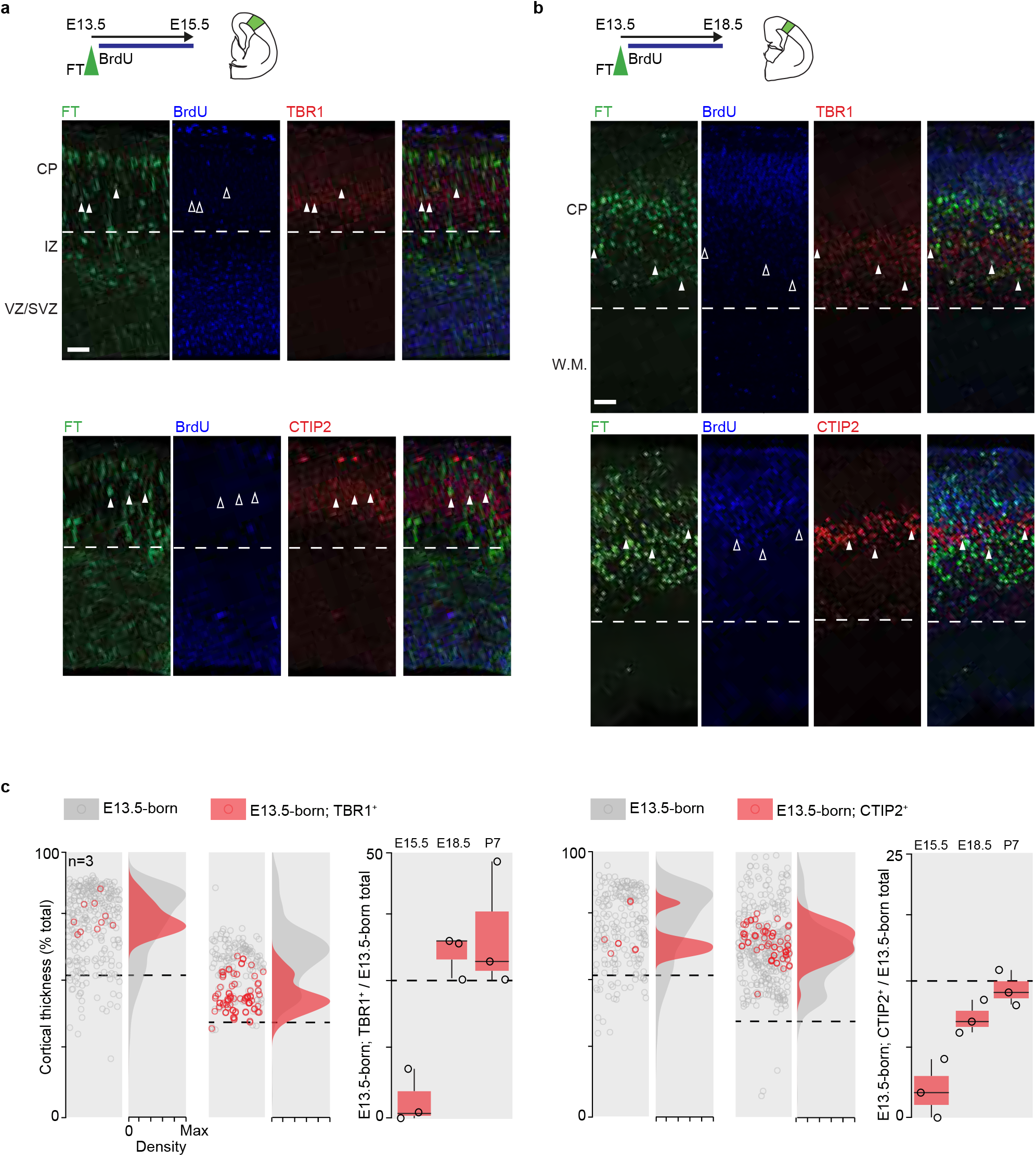
Simultaneously early born neurons progressively acquire molecular diversity during differentiation. **a**, Immunostaining of E13.5 AP-born neurons with TBR1 and CTIP2 antibodies at E15.5. **b**, Immunostaining of E13.5 AP-born neurons with TBR1 and CTIP2 antibodies at E18.5. **c**, The trend of E13.5 AP-born neurons expressing TBR1 and CTIP2 display a progressive acquisition of molecular diversity during differentiation. Scale bars: 50μm (a, b). CP – Cortical Plate, E – embryonic day, FT – FlashTag, IZ – Intermediate Zone, L – Layer, P – Postnatal day, SVZ – Subventricular Zone, VZ – Ventricular Zone, WM – White matter.

**Figure S5.**
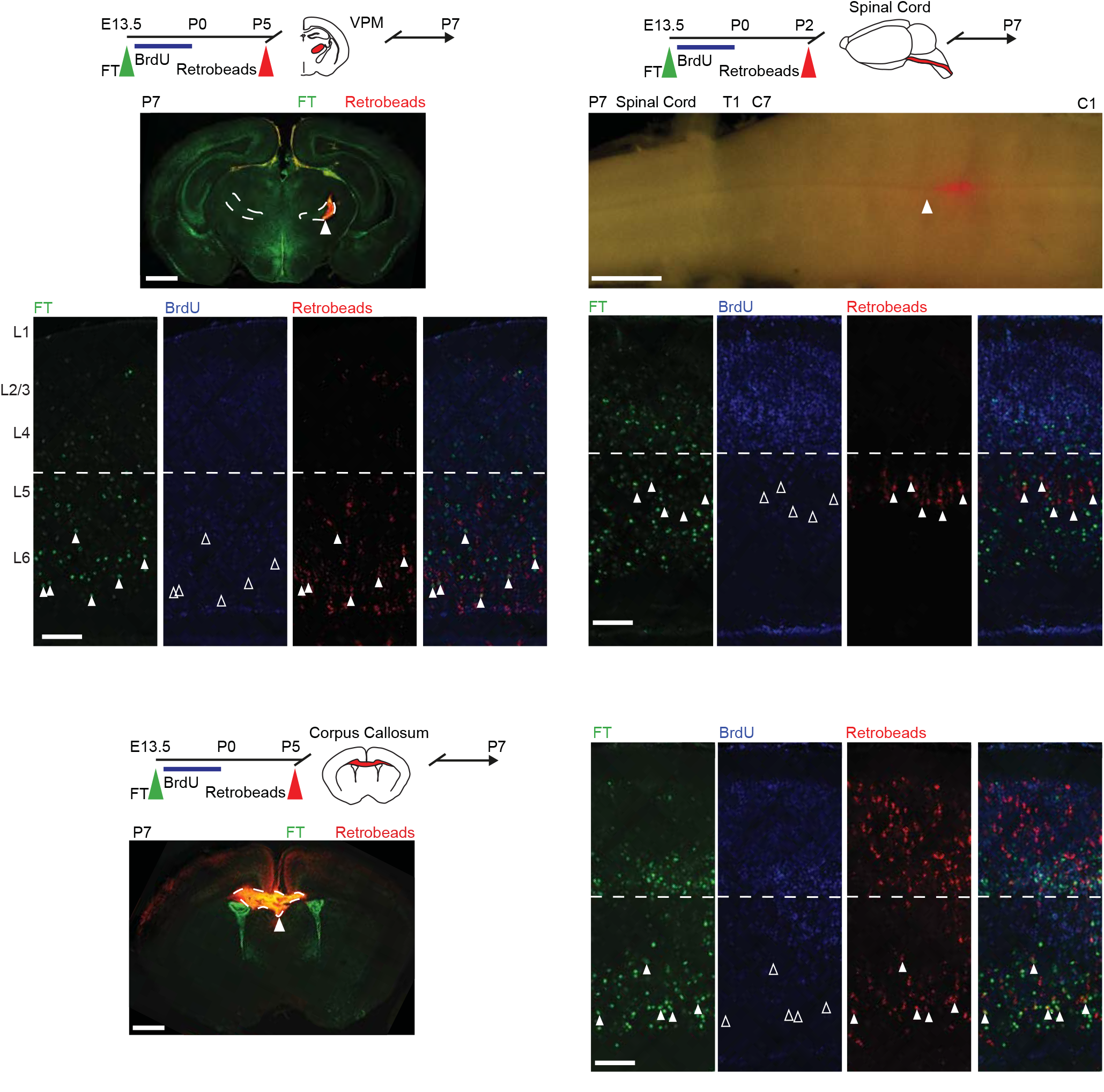
Simultaneously early born neurons diverse laminar distribution and connectivity. Retrograde labeling of E13.5 simultaneously-born neurons reveals a diversity of projections to the ventroposterior medial nucleus of the thalamus (VPM), spinal cord and corpus callosum. Scale bars: 200μm (high mag), 1mm (low mag). E – embryonic day, FT – FlashTag, L – Layer, P – Postnatal day, S1 – primary somatosensory area, VPM – Ventro Posterior Medial Thalamic nucleus.

